# Characterization of plasma circulating small extracellular vesicles in patients with metastatic solid tumors and newly diagnosed brain metastasis

**DOI:** 10.1101/2021.11.18.469115

**Authors:** Alberto Carretero-González, Sara Sánchez-Redondo, Luis Manso Sánchez, Eva Ciruelos Gil, Daniel Castellano, Marta Hergueta-Redondo, Guillermo de Velasco, Héctor Peinado

## Abstract

**Purpose:** Nearly 40% of the advanced cancer patients will present brain metastases during the course of their disease, with a 2-year life expectancy of less than 10%. Immune system impairment, including the modulation of both STAT3 and PD-L1, is one of the hallmarks of brain metastases. Liquid biopsy could offer several advantages in brain metastases management, such as the possibility of non-invasive dynamic monitoring. Extracellular vesicles (EVs) have been recently proposed as novel biomarkers especially useful in liquid biopsy due to their secretion in biofluids and their role in cell communication during tumor progression.

**Materials and Methods:** The main aim of this work was to characterize the size and protein cargo of plasma circulating EVs in patients with solid tumors and their correlation with newly diagnosed brain metastases, in addition to their association with other relevant clinical variables.

**Results:** We analyzed circulating EVs in the plasma of 123 patients: 42 patients with brain metastases, 50 without brain metastases and 31 healthy controls. Patients with newly diagnosed brain metastases had a lower number of circulating EVs in the plasma and a higher protein concentration in small EVs (sEVs) compared to patients without brain metastases and healthy controls. Interestingly, melanoma patients with brain metastases presented decreased STAT3 activation and increased PD-L1 levels in circulating sEVs compared to patients without central nervous system metastases.

**Conclusions:** Decreased STAT3 activation and increased PD-L1 in plasma circulating sEVs identify melanoma patients with brain metastasis.

**Statement of translational relevance:** Brain metastases are critical for outcomes and quality of life in almost 50% of oncological patients, generally associated with a poor short-term prognosis. Early or preventive diagnosis of this complication represents an unmet need. There is a necessity of discovering new biomarkers that could aid to predict disease outcome.

In this study, we analyzed plasma circulating extracellular vesicles (EVs) from a cohort of 92 patients with different solid tumors (lung, breast, kidney cancer and melanoma) and found that newly diagnosed patients with brain metastases presented lower number of circulating particles and a higher protein concentration in small extracellular vesicles (sEVs) compared to patients without brain metastases and healthy controls. Out of all groups analyzed, melanoma patients with brain metastases presented decreased STAT3 activation and increased PD-L1 levels in circulating sEVs compared to patients without central nervous system metastases.

The data presented in this work suggest that circulating sEVs may represent the immunosuppressive status of newly diagnosed brain metastases characterized by the reduced phospho-STAT3 (pSTAT3) and increased PD-L1, although the origin of these molecules found in circulating sEVs remains to be uncovered.

## Introduction

One out of 10 patients with solid tumors will develop brain metastases ^1^, 40% of patients with metastatic tumors ^1,2^, during the course of their disease. The main tumor types with higher incidence of brain metastasis are lung cancer (40-50% of cases), breast cancer (15-25%) and melanoma (5-20%) ^2^. Despite the current multimodal strategies, central nervous system (CNS) metastases are associated with poor outcomes (less than 10% of the patients are alive after 2 years from the diagnosis) and detrimental quality of life of the patients ^3^. Differential factors involved in brain metastases dissemination could justify the failure of systemic therapies such as the presence of the blood-brain barrier (BBB) or evolutionary divergences found in cell clones localized in the CNS compared to extracranial metastases ^4,5^.

The induction of an immunosuppressive state is another factor involved in CNS metastatic spread ^6^. Out of many factors involved, the signal transducer and activator of transcription 3 (STAT3) is particularly relevant ^7^. Its activity has been shown to increase in brain metastases compared to primary tumors in melanoma, lung or breast cancer ^7–10^. Moreover, STAT3-activated reactive astrocytes within brain metastases have been shown to inhibit CD8^+^ T lymphocyte activation through the expression of programmed death-ligand 1 (PD-L1) and the secretion of other immunosuppressive molecules, favoring tumor growth ^11^. Indeed, STAT3 is involved in immune cell evasion aiding tumor progression and metastasis ^12^. Similarly, PD-L1 expression has also been reported in different brain stromal cells with immunosuppressive consequences, such as reactive astrocytes ^11^, microglial cells ^13^ or infiltrating peripheral neutrophils ^14^.

Small extracellular vesicles (sEVs) represent a subtype of extracellular vesicles (EVs) ^15,16^. In malignant diseases, EVs play an important role as intercellular communicators in the establishment of the pre-metastatic niche ^17,18^ and brain metastasis ^17,19–21^. EVs represent one of the last elements to be included in liquid biopsy studies ^22,23^; plasma samples are the main source analyzed so far. In recent years, secretion of PD-L1 in EVs has been pointed out as a potential mechanism of resistance to the treatment with immune checkpoint inhibitors ^24,25^. For example, exogenous administration of tumor cell-derived EVs, expressing PD-L1 on their surface, increased the ability of metastatic dissemination and primary tumor growth ^26^. Plasma circulating PD-L1 from plasma decreased the activation of CD8^+^ lymphocytes T ^27^. On the other hand, the secretion of STAT3 in plasma-circulating sEVs has not been reported. Considering the role of both PD-L1 and phospho-STAT3 (pSTAT3) in the biology of brain metastases, we wanted to characterize the presence of these molecules in plasma circulating sEVs in patients with newly diagnosed brain metastases to identify their relevance as minimally invasive biomarkers for the CNS metastatic dissemination.

## Materials and methods

### Study design, population and sample collection

Observational, single academic center study. Plasma samples were collected prospectively from February 2017 to December 2020.

Two different sets were considered

a. Patients diagnosed with metastatic solid tumors (lung, breast, kidney cancer or melanoma, the most frequent sources known to spread to the CNS). They were prospectively included with either confirmed progressive disease state or a recent diagnosis of their metastatic tumor. Patients were not receiving any potential systemic therapy interfering with the analyses. The assessment of the brain metastatic status was performed according to usual practice, using standard imaging tests such as brain computed tomography (CT) or brain magnetic resonance imaging (MRI). Those patients with brain metastases had to present a recent (*de novo*) diagnosis of their CNS involvement at the time of sample collection; patients with previous history of brain dissemination were excluded. Patients considered to have no brain metastases were also assessed by radiological tests in order to confirm their status.
b. Healthy controls. Healthy volunteers and patients with a previous diagnosis of any solid tumor radically treated and without evidence of local or distant relapse during at least 5 years of follow-up (cured controls) were included.

The clinical/pathological variables collected are detailed in **Supplementary Table 1**. The protocol of this study was reviewed and approved by the Committees for Ethical Research (identification number 17/052), as indicated by the Spanish legislation. All samples were obtained after signing informed consent. The study was conducted in accordance with the ethical principles of the Declaration of Helsinki and Good Clinical Practice guidelines.

Once the inclusion criteria mentioned above were confirmed, blood samples (approximately 10 ml) were collected in EDTA tubes and centrifuged for 10 min at 500 x g at room temperature using a swing out bucket rotor. The plasma fractions (supernatant) were poured into collection tubes and immediately frozen at −80°C.

### sEV isolation

Purification of sEVs from plasma was performed after thawing the samples at 37°C for 3–5 min. First, samples were centrifuged at 3,000 × g for 20min, followed by further centrifugation of the supernatants at 12,000 × g for 20 min. The sEVs were subsequently harvested by centrifugation of the supernatant fraction at 100,000 × g for 70 min. The sEV pellet was resuspended in 3.5 ml of PBS and collected by a second ultracentrifugation at 100,000 × g for 70 min. All centrifugations were performed at 10°C using a Beckman OptimaX100 centrifuge with a Beckman 70.1Ti rotor.

sEVS were resuspended in 100 µl PBS and the protein content was measured by bicinchoninic acid (BCA) assay (Pierce). Particle number was measured from an aliquot of 1 μl of plasma sEVs diluted in 1 ml of PBS using NTA (NanoSight; Malvern) equipped with a violet laser (405 nm).

### Immunoblotting

sEVs were lysed in Laemmli buffer at 95°C for 5 min, and 10 µg of the protein extracts were resolved by SDS-PAGE and probed using antibodies against STAT3 (1:1,000, #9139S; Cell Signaling), Phospho-STAT3 (1:2,000, #9145S; Cell Signaling), PD-L1 (1:1,000, #13684S; Cell Signaling), β-actin (1:10,000, #A5441; Sigma-Aldrich) and Alix (1:1,000, #2171S; Cell Signaling). Peroxidase-conjugated AffinityPure donkey anti-rabbit or anti-mouse (1:5,000; Jackson ImmunoResearch) were used as secondary antibodies. Signal was detected using ECL Western Blotting Substrate kit (GE Healthcare). The intensity of the immunoreactive bands was quantified by densitometry using ImageJ software (National Institutes of Health).

### Endpoints

We aim to characterize plasma-circulating sEVs from patients with recently diagnosed brain metastases and compare them to those from metastatic patients without CNS involvement and healthy controls. Specifically, particle number in plasma, protein concentration and expression of pSTAT3, STAT3 and PD-L1 by Western blot in plasma-circulating sEVs were quantified and correlated with clinically relevant characteristics, especially the recent diagnosis of brain metastases. The type of response achieved with the immediate treatment received after plasma sample collection and overall survival of patients were also assessed and correlated with the parameters analyzed in plasma-circulating sEVs.

### Statistical analysis

GraphPad Prism (version 8.1.1) was used for all statistical analyses. Baseline patient characteristics were compared between the study groups (concretely, patients with *vs* without brain metastases) using *t* tests, Mann-Whitney test (quantitative variables) or two-tailed Fisher’s exact test and Chi-square test (qualitative variables). The differences for the parameters analyzed were assessed by parametric tests (one-way ANOVA test, unpaired Student’s test) and nonparametric tests (Kruskal-Wallis test, Mann-Whitney test). Overall survival (OS) analyses were performed using the Kaplan-Meier method; differences were evaluated using the Log-Rank test. Controlled subgroup analyses were performed to assess the influence of potential confounding factors. Statistical significance was established for a *p* value < 0.05.

## Results

### Analysis of circulating EVs in plasma

A total of 123 patients who met the pre-specified inclusion criteria were finally included: 42 metastatic patients with brain metastases, 50 patients without brain metastases and 31 healthy controls **(Table 1)**. Specifically for metastatic patients with or without brain metastasis, no statistically significant differences were found in relation to the number or type of previous systemic therapies received between these groups. See **Supplementary Tables 2 and 3** for further details. Metastatic patients with CNS involvement had a shorter OS (time from diagnosis of metastatic disease to death due to any cause) (median OS 25.20 months) compared to those without brain metastases (median OS 55.80 months) (Hazard ratio 1.99; 95% confidence interval 1.11-3.57) **(Figure 1A)**. A consistent trend was observed in all histological types considered, with a worse outcome for those patients with CNS dissemination **(Supplementary Figures 1A-1D)**.

**Table 1.** Clinical characteristics of the patients included in the study at the time of sample collection. General characteristics of the main groups of patients included in the study are shown. CNS: Central nervous system.

|  | Metastatic patients with CNS disease (n=42) |  | Metastatic patients without CNS disease (n=50) |  | Healthy/cured controls (n=31) |  |
| --- | --- | --- | --- | --- | --- | --- |
| **Histological type of the primary tumor** | **n** | **%** | **n** | **%** | **n** | **%** |
| Lung cancer | 13 | 31 | 12 | 24 | — | — |
| Breast cancer | 14 | 33 | 21 | 42 | — | — |
| Kidney cancer | 4 | 10 | 10 | 20 | — | — |
| Melanoma | 11 | 26 | 7 | 14 | — | — |
| **Number of previous systemic therapies received for metastatic disease** |  |  |  |  |  |  |
| 0 | 18 | 43 | 28 | 56 | — | — |
| 1 or more | 24 | 57 | 22 | 44 | — | — |
| **Type of systemic therapies received for metastatic disease** |  |  |  |  |  |  |
| Chemotherapy | 17 | 40 | 11 | 22 | — | — |
| Immune checkpoint inhibitors | 7 | 17 | 4 | 8 | — | — |
| Targeted therapy (including hormone therapy) | 24 | 57 | 33 | 66 | — | — |
| No systemic treatment | 18 | 43 | 28 | 56 | — | — |
| **Previous oncological history** |  |  |  |  |  |  |
| No oncological history | — | — | — | — | 10 | 32 |
| Breast cancer | — | — | — | — | 9 | 29 |
| Testicular tumor (germ cell) | — | — | — | — | 12 | 39 |
| **Time elapsed since curative treatment** |  |  |  |  |  |  |
| 5-7 years | — | — | — | — | 13 | 42 |
| 7-10 years | — | — | — | — | 5 | 16 |
| >10 years | — | — | — | — | 3 | 10 |
| **Previous adjuvant/neoadjuvant systemic treatment** |  |  |  |  |  |  |
| Chemotherapy | — | — | — | — | 15 | 49 |
| No systemic treatment | — | — | — | — | 6 | 19 |

**Figure 1.**
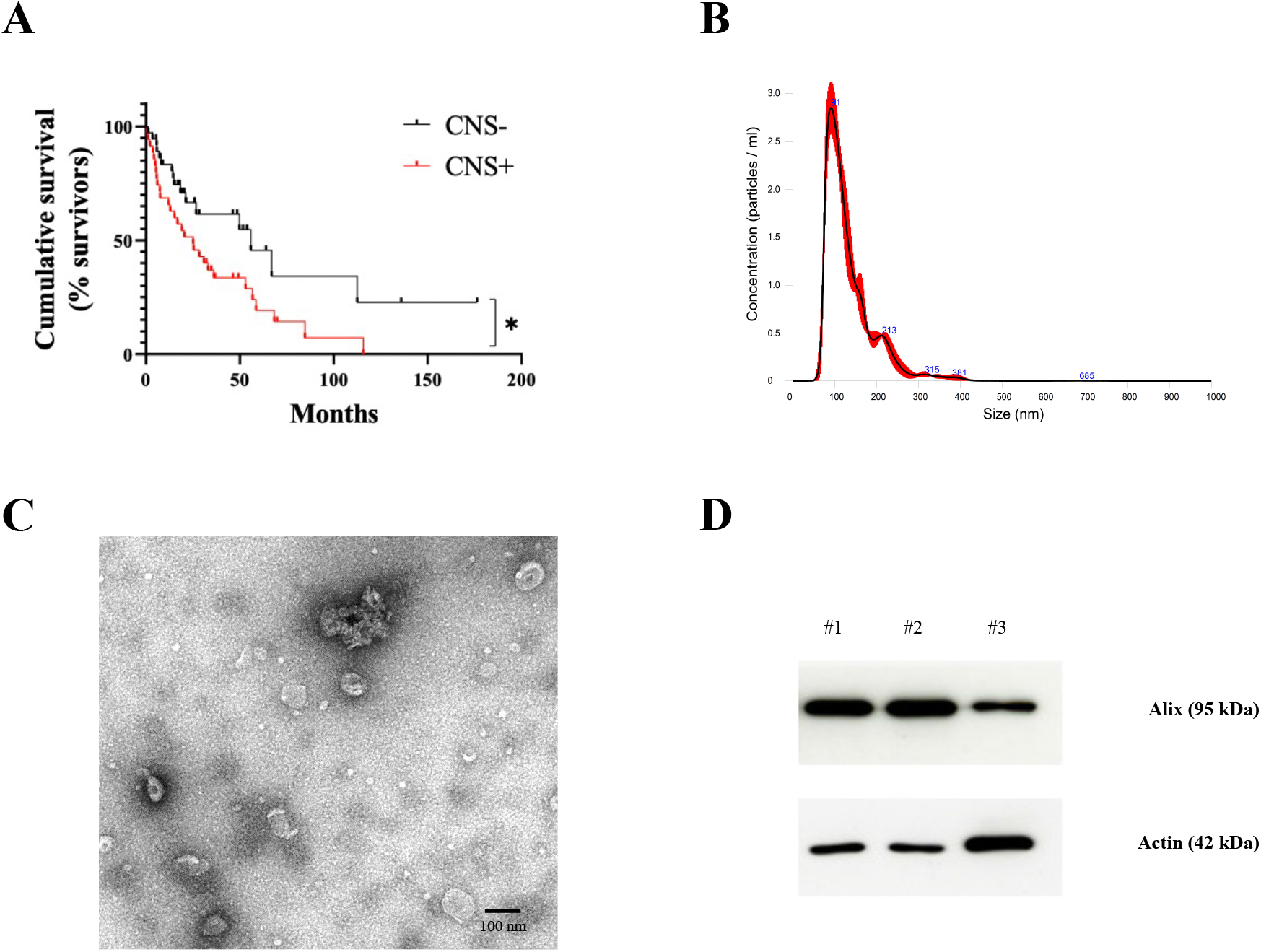
Overall survival analysis of patients included in the study and characterization of plasma-circulating sEVs. **A**. Survival analysis and graphical representation using Kaplan-Meier curves showing the cumulative survival probability in the study population according to the absence (CNS-) or presence (CNS+) of central nervous system (CNS) metastases. * *p* value 0.02. Differences were assessed using the Log-Rank test. **B**. Representative image of the particle content (x10^8^) by NTA analysis of a plasma sample from a melanoma patient. **C**. Representative electron microscopy imaging of sEVs from the same patient’s plasma. **D**. Western blot analysis of the known exosome marker Alix in sEVs isolated from the plasma of three different patients. Actin was used as loading control.

In order to characterize plasma circulating EVs, we measured their number using nanoparticle track analysis (NTA) **(Figure 1B)**. sEV integrity was confirmed by electron microscopy **(Figure 1C)**. We verified the enrichment of typical sEV markers such as Alix in our preparations ^28^ **(Figure 1D)**.

The analysis of particle number in plasma samples showed that lung, breast and kidney cancer patients with CNS metastases presented a decreased number of particles compared to those patients without brain metastasis **(Figures 2A-2C)**. In melanoma patients we did not find significative differences among groups **(Figure 2D)**. On the other hand, CNS metastasis was associated with an increased protein concentration in plasma-circulating sEVs from lung cancer and breast cancer patients compared to those without brain metastases **(Figures 2E-2F)**. Although not statistically significant, there was also a trend for an increased protein concentration in the circulating sEVs of kidney cancer patients with CNS metastases **(Figure 2G)**. Regarding melanoma patients, no differences were found in relation to sEV protein concentration according to the CNS status **(Figure 2H)**. See **Supplementary Tables 4 and 5** for further details.

**Figure 2.**
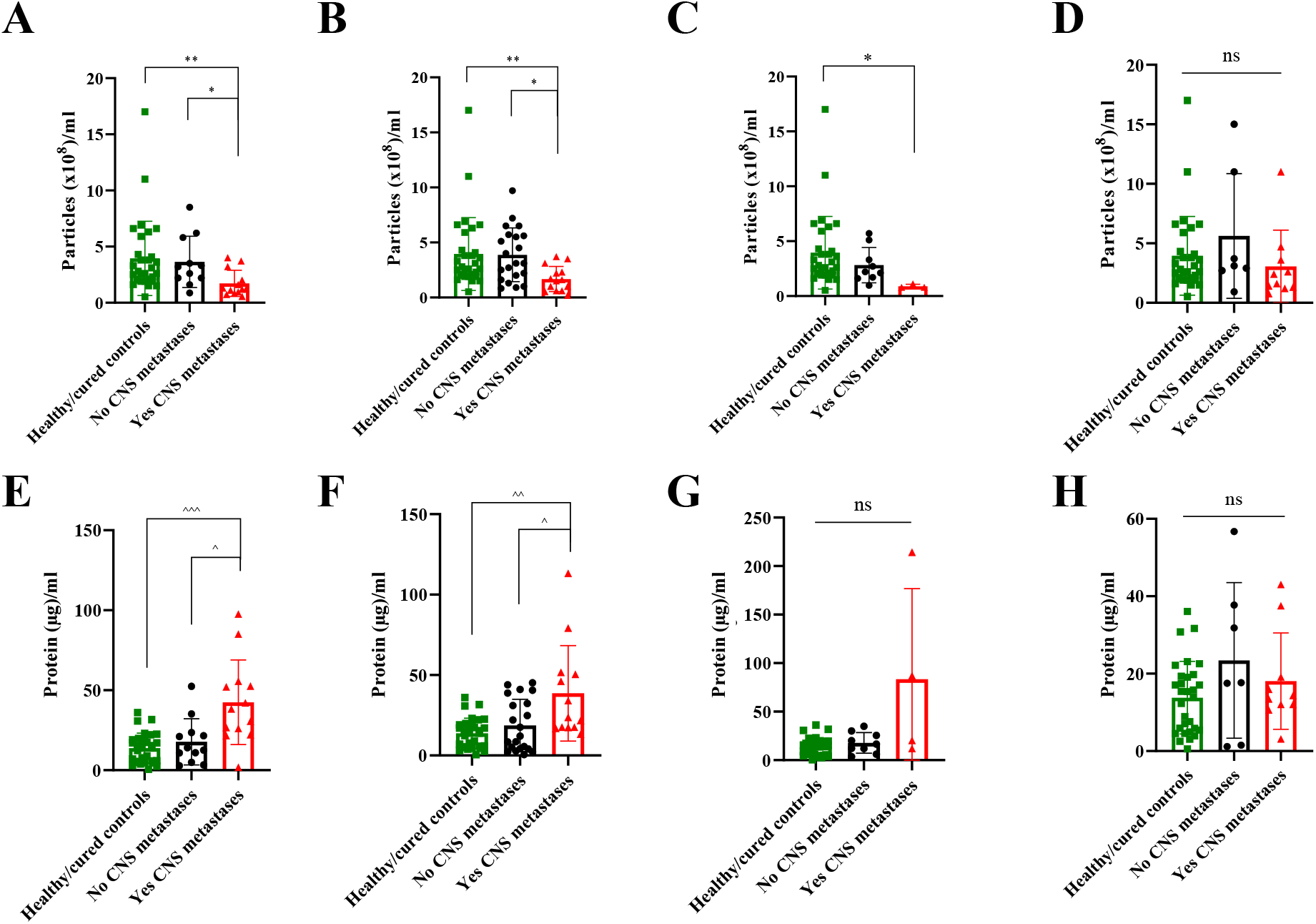
Quantitative analysis of plasma-circulating sEVs from patients included in the study according to their central nervous system (CNS) metastases status (including healthy/cured controls). **A**. Particles/ml in lung cancer. **B**. Particles/ml in breast cancer. **C**. Particles/ml in kidney cancer. **D**. Particles/ml in melanoma. **E**. Proteins/ml in lung cancer. **F**. Proteins/ml in breast cancer. **G**. Proteins/ml in kidney cancer. **H**. Proteins/ml in melanoma. * *p* value < 0.05. ** *p* value < 0.01. ^ *p* value < 0.05. ^^ *p* value 0.003. ^^^ *p* value 0.0003. ns: not significant.

Analysis of correlation of protein concentration with patient survival showed that a high protein concentration in plasma-circulating sEVs is associated with a more aggressive clinical course and a reduced overall survival in lung cancer and melanoma patients **(Figures 3A and 3B)**. No differences were found according to the immunohistochemical subtypes for breast cancer and kidney cancer patients or according to absence/presence of previous oncological history in the healthy controls group (data not shown).Specifically in breast cancer, a higher protein concentration was found in patients with a more aggressive clinical course of their disease (tumor progression at multiple levels, including CNS and other systemic locations) compared to patients with CNS progression only **(Figure 3C)**.

**Figure 3.**
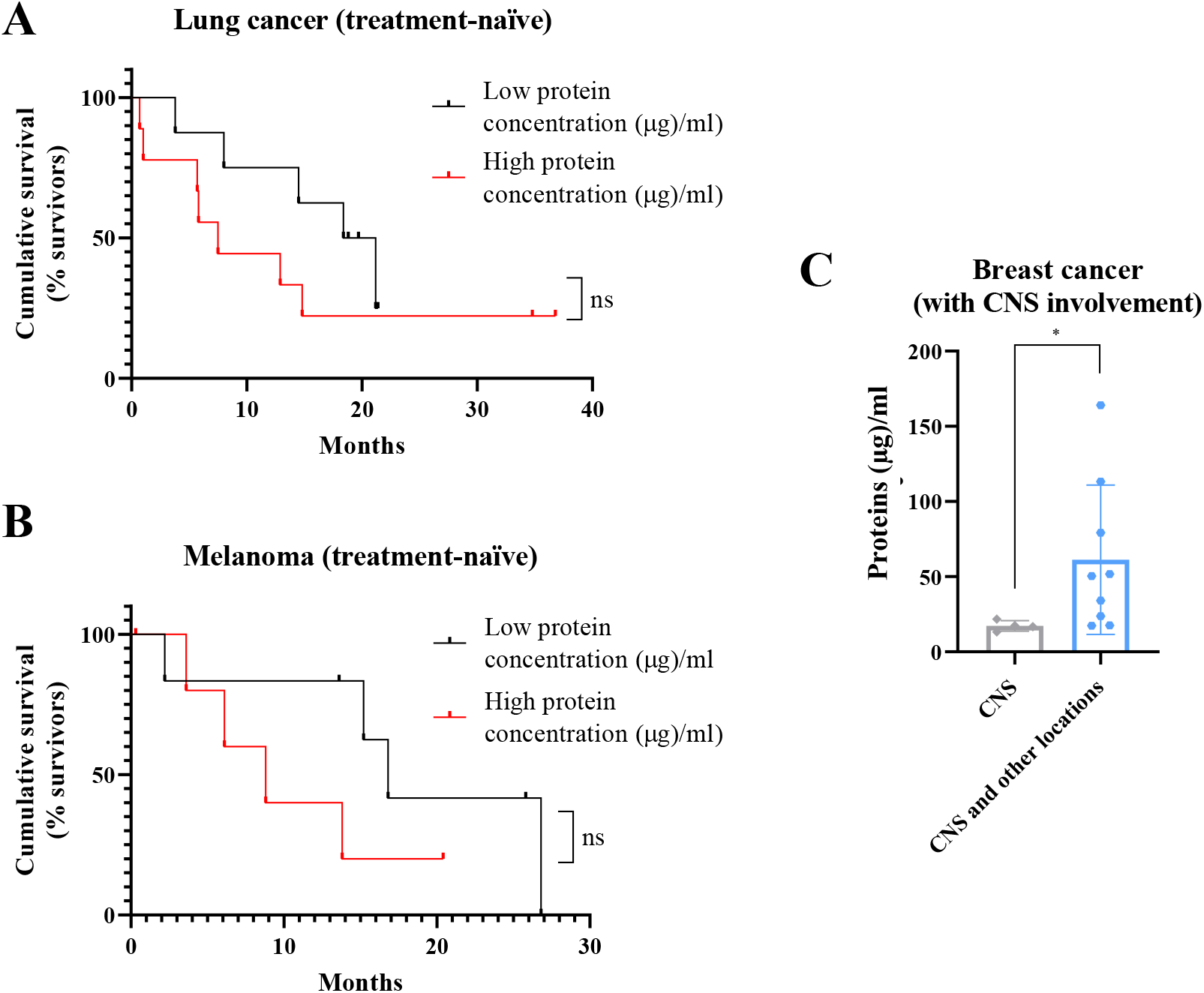
Complementary studies for total protein concentration in patients plasma-circulating sEVs showing a correlation between a high protein amount and a worse prognosis. **A**. Survival analysis showing the cumulative survival probability in patients with previously untreated metastatic lung cancer according to the protein concentration in plasma-circulating sEVs (taking the median value of the group as reference). **B**. Survival analysis showing the cumulative survival probability in patients with previously untreated metastatic melanoma according to the protein concentration in plasma-circulating sEVs (taking the median value of the group as reference). Differences were assessed using the Log-Rank test. **C**. Analysis of protein concentration in circulating sEVs regarding the type of progression experienced in patients with metastatic breast cancer and known central nervous system (CNS) involvement: at CNS only or at CNS and other locations. * *p* value 0.02. ns: not significant.

### Characterization of PD-L1 and pSTAT3 expression by Western blot in plasma-circulating sEVs

Due to the relevance of STAT3 and PD-L1 in brain metastasis progression ^7–11,13,14^, we wondered if the analysis of these molecules in circulating sEVs could be modulated along metastatic progression. The comparative analysis of PD-L1 expression in plasma-circulating sEVs from patients with lung, breast o kidney cancer showed no differences regarding the CNS metastatic status **(Figure 4A)**. Regarding the analysis of STAT3, a double band at the expected weight of STAT3 was observed in most of the patients analyzed, fact that could be explained by the simultaneous presence of two isoforms (STAT3α and STAT3β as a consequence of alternative splicing) ^29^ **(Supplementary Figures 2A, 2B and 3A)**. However, unlike in melanoma patients (see below), a double band was not clearly observed in the pSTAT3 analysis for lung, breast and kidney cancer patients **(Supplementary Figures 2A, 2B and 3A)**; thus, the quantification of pSTAT3 expression in these cases was performed considering exclusively the band observed at the expected weight for this protein ^29^. Patients with metastatic breast cancer and brain metastases presented an increased STAT3 activation in their plasma-circulating sEVs compared to patients without CNS involvement; in this regard, no differences were found for lung or kidney cancer patients **(Figure 4B)**.

**Figure 4.**
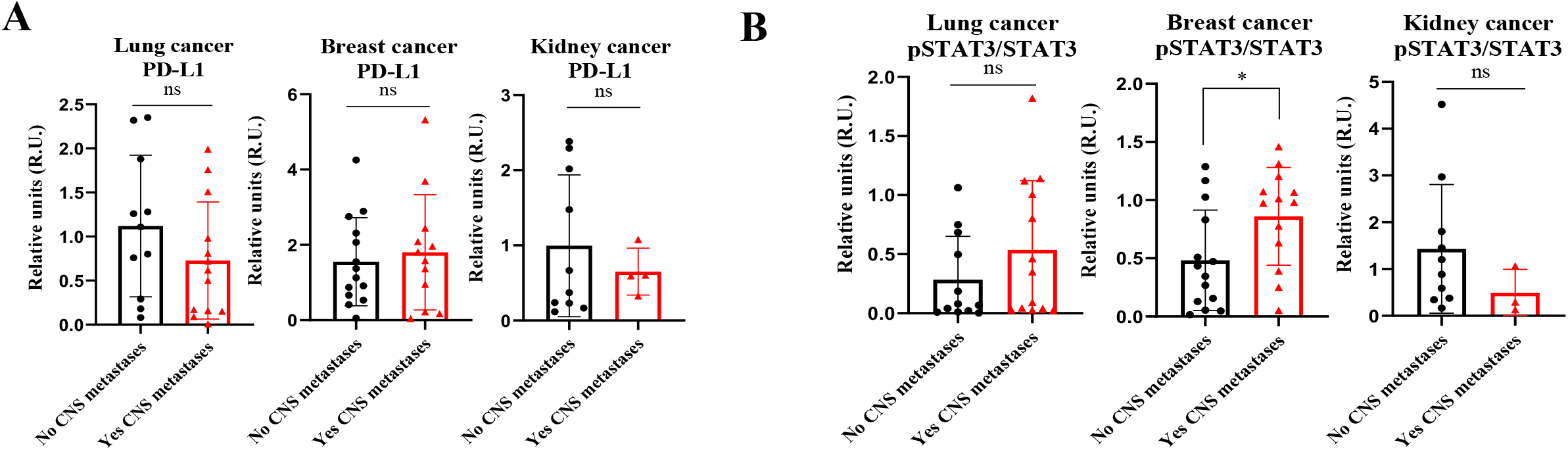
Analysis of PD-L1 and pSTAT3/STAT3 expression by Western blot in plasma-circulating sEVs from patients with lung, breast or kidney cancer. **A**. Quantification of PD-L1 expression levels and statistical analysis of samples obtained. The data obtained for PD-L1 in densitometry were normalized to Ponceau values. **B**. Quantification of pSTAT3/STAT3 expression levels and statistical analysis of samples obtained. * *p* value 0.03. ns: not significant.

Interestingly, the analysis of PD-L1 levels in melanoma patients showed an increased expression of this protein in circulating sEVs from the patients with a recent diagnosis of brain metastases compared to those without known CNS dissemination **(Figures 5A and 5B)**. In contrast to the rest of the patients considered in this series, melanoma cases without known brain metastasis presented a double band in the pSTAT3 analysis; both pSTAT3α (higher molecular weight) and pSTATβ (lower molecular weight) expressions were retained **(Figure 5A)**. In this tumor subtype, we unexpectedly observed that pSTAT3α showed a strong dephosphorylation in patients with brain metastasis **(Figures 5A and 5C)**.

**Figure 5.**
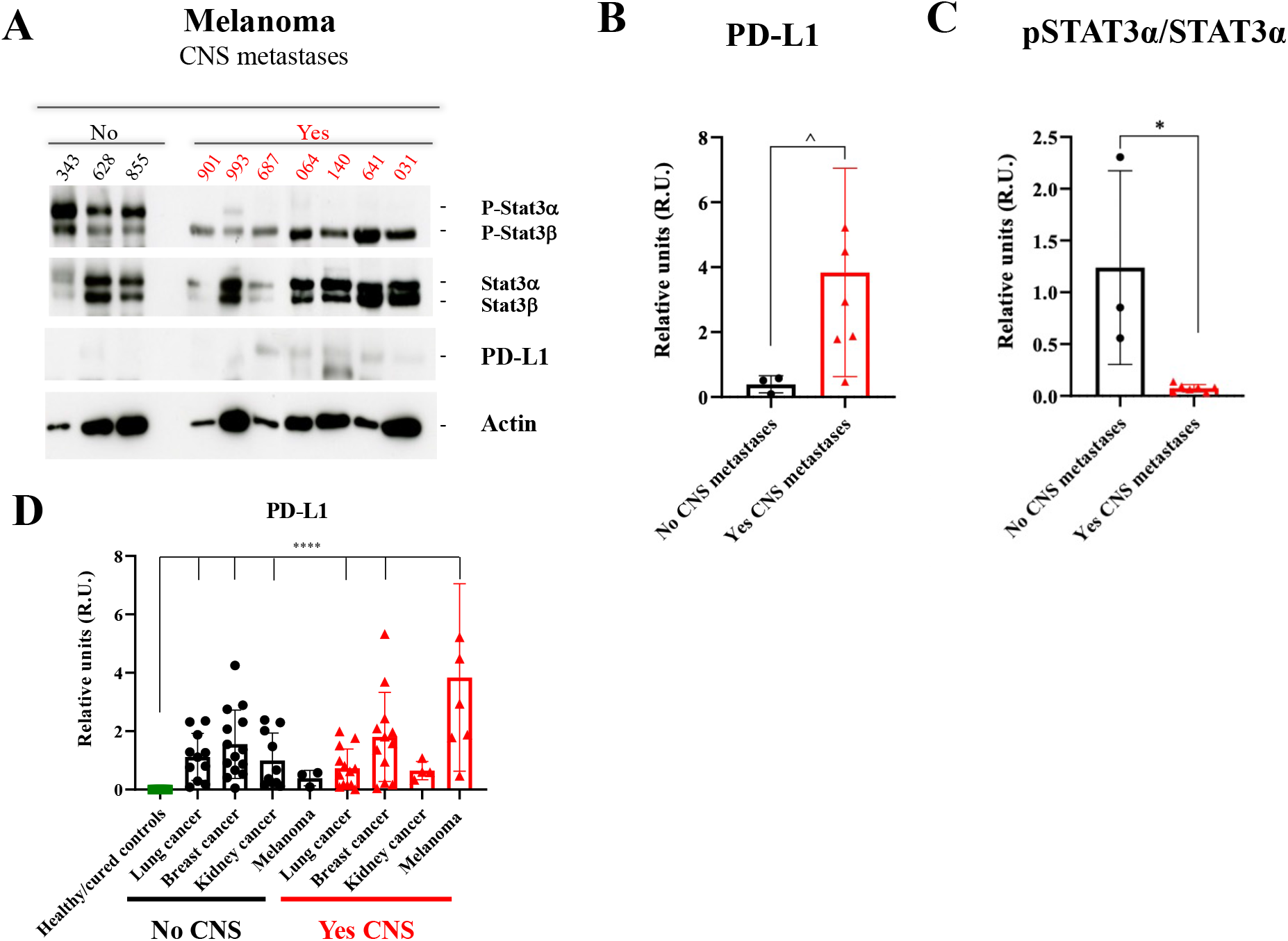
Study of pSTAT3/STAT3 and PD-L1 expression by Western blot in plasma-circulating sEVs from patients with melanoma. **A**. Western blot image of the expression of different proteins (indicated on the right) in sEVs from melanoma patients according to the absence (No) or presence (Yes) of central nervous system (CNS) metastases. **B**. Quantification of PD-L1 expression levels and statistical analysis of samples considered in **A**. The data obtained for PD-L1 in densitometry were normalized to Ponceau values. **C**. Quantification of pSTAT3α/STAT3α band expression levels and statistical analysis of samples considered in **A. D**. Quantification of PD-L1 expression levels and statistical analysis of the whole population of this study, including the healthy/cured controls group. The data obtained for PD-L1 in densitometry were normalized to Ponceau values. No CNS: Absence of CNS metastases. Yes CNS: Presence of CNS metastases. ^ *p* value 0.03. * *p* value 0.02. **** *p* value < 0.0001.

Strikingly, some patients with low pSTAT3/STAT3 sEVs expression, regardless of their tumor subtype or CNS status, were repeatedly associated with an increased PD-L1 level **(Supplementary Figures 2A, 2B and 3A)**. Of note, sEVs purified from the plasma of healthy patients showed a complete absence of PD-L1 expression and very low levels of pSTAT3 **(Supplementary Figure 3B)**.

In the global analysis of PD-L1 expression **(Figure 5D)**, a statistically significant absence of this protein was found in the healthy controls group; this quantification also showed that there is a significant increase of PD-L1 in circulating sEVs regardless of the patients group analyzed except in melanoma patients with no dissemination to the SNC and kidney cancer patients with dissemination to the SNC. These data suggest that patients with no active cancer have undetectable levels of PD-L1 in plasma circulating sEVs.

## Discussion

Tests using biological fluids for detecting tumor material are a non-invasive approach complementary or even alternative to tissue biopsies. Circulating tumor cells (CTCs) and circulating tumor DNA (ctDNA) are considered the cornerstones of liquid biopsy-based diagnosis ^30^. However, in addition to CTCs and ctDNA, the analysis of tumor-secreted EVs have been of increasing interest in recent years. Although there remains debate about the benefits of circulating EV compared to other circulating biomarkers, there have been several studies demonstrating their utility and benefits in diagnostics using specific biomarkers including DNA, RNA and protein biomarkers ^31,32^.

Several studies support an association between an increased number of plasma particles and a worse outcome or aggressive features in oncological patients with colon ^33^ or head and neck cancer ^34^, among others. However, other studies have found no differences in the number of circulating EVs analyzed in plasma or other biological fluids from melanoma patients according to the tumor stage or the extent of loco-regional lymph node involvement ^35,36^, similar to the results here presented for this specific subtype.

These differences could probably by explained by the heterogeneity of the series considered, as well as the absence of a standardized methodology for sample handling or the divergent biology of the different tumor types ^35^. In our series, we observed that the number of circulating particles was reduced in breast, lung and kidney cancer patients with brain metastases. Although the reasons behind these changes are hard to interpret, we observed that, concomitantly with these changes, breast and lung cancer patients with brain metastases showed increased amount of protein associated to circulating sEVs. Although scarce, data from different studies regarding the value of total protein concentration in circulating sEVs point to an association between a higher protein concentration in circulating sEVs and a worse clinical course ^35–38^. In our series, the analysis of the correlation between the total protein content in sEVs and patient survival suggests a potential prognostic value. Measurement of protein concentration in sEVs could be useful for the assessment of tumor dissemination to the CNS; nevertheless, since we did not reach statistical significance in these cohorts, analyses in larger cohorts are needed. Similarly, since total protein in sEVs may indicate systemic changes (e.g. EVs derived from circulating platelets, immune cells, etc…) the source of this material as well as the biological relevance needs to be solved.

Tumor-secreted EVs have the intrinsic ability to breach biological barriers such as the BBB ^39^. Brain metastases, result from the dissemination of tumor cells to the brain, most commonly from lung cancer, melanoma, and breast cancer ^1,2^. EVs contribute to different stages of brain metastasis increasing BBB permeability ^21,39^, reprograming of brain metastatic niches ^20,40^ and increasing brain metastatic organotropism of tumor cells ^18,19^. Overall, these studies demonstrate that EVs can drive bidirectional cross talk between tumor cells and their microenvironment promoting metastasis formation in the brain. However, the analysis of EVs in clinical samples is unreported, to the best of our knowledge, this study constitutes the first approach to characterize the potential clinical value of a non-invasive technique, liquid biopsy using plasma-circulating sEVs, in patients with different solid tumors and recently diagnosed brain metastases.

To perform these analyses the selection of plasma samples was carried out following well-established criteria to ensure the homogeneity of the selected group and to limit possible confounding factors common in clinical practice. It is noteworthy that none of the oncological patients were under active systemic treatment potentially influencing the results at the time of sample collection; in addition, the group of patients with CNS involvement consisted of patients with no previously known brain disease (*de novo* diagnosis). This series of patients consisted of three distinct groups of subjects. The healthy controls, when compared with the other two oncological groups, allowed us to characterize features inherent to any oncological disease. Specifically, the presence of two defined groups of metastatic patients, which differed only in the presence of CNS involvement, allowed to establish associations between the results obtained and the metastatic brain dissemination process.

Although no specific studies have been performed in patients with brain metastatic dissemination, there are data reporting an association between the EV expression of PD-L1 and pSTAT3 and a more advanced oncological disease, more aggressive clinical course and, even, mechanisms of treatment resistance ^24,25,41^.

STAT3α, the predominant STAT3 splice form in several cell types, typically shows rapid phosphorylation and nuclear translocation following cytokine stimulation, but their differential role is still a matter of discussion ^42^. We have observed that plasma sEVs from melanoma patients with brain metastases showed reduced pSTAT3α levels in addition to an increased PD-L1 expression compared to those from melanoma patients without CNS dissemination, characterized by higher levels of pSTAT3 together with the absence of PD-L1. The fact that pSTAT3α disappears from circulating sEVs suggest that it could be executing its actions intra-nuclearly in immune and/or tumor cells preventing its shedding in plasma-circulating sEVs. Indeed, activated STAT3 in immune cells results in inhibition of immune mediators and promotion of immunosuppressive factors ^43^ as well as melanoma metastatic behavior ^44^. Moreover, STAT3 inhibition reduced melanoma metastasis to brain ^45^ and has recently been hypothesized to be considered a potential target for immunotherapy ^46^. Interestingly, a comparative study between lung cancer patients with and without CNS metastases also showed a higher concentration of PD-L1^+^ myeloid cells, immunosuppressive myeloid cells and regulatory T lymphocytes in the peripheral blood of patients with brain involvement, all of them considered as systemic immunosuppression markers ^47^. In summary, we observed a decrease of pSTAT3α levels and increased PD-L1 expression in patients with CNS dissemination suggesting that these changes are a consequence of the systemic immunosuppression observed in melanoma brain metastatic patients. These data suggest that the analysis of these molecules in plasma-circulating sEVs in melanoma patients with brain metastasis undergoing immunotherapy or in the combination with STAT3 inhibitors could be useful to monitor treatment response.

On the other hand, no significant differences were found for the expression of PD-L1 and pSTAT3 in plasma-circulating sEVs from lung, breast or kidney cancer patients depending on the presence or absence of newly diagnosed brain metastases, suggesting that these diseases depend on other molecular pathways. However, complementary analyses in some of these tumor types suggested an association between a pattern consisting of a reduced pSTAT3 or an increased PD-L1 expression (similar to that of melanoma patients with brain metastases) and a more aggressive clinical course.

Noteworthy, we observed an inverse correlation between the pSTAT3 and PD-L1 expressions in plasma-circulating sEVs in most of the tumor types considered. Further mechanistic studies are warranted to explain these findings highlighting the relationship between EV expression and cellular activation of these proteins. Our findings also support the potential value of sEV-derived PD-L1 as a diagnostic marker of active oncological disease in several types independently of brain metastasis (e.g. breast cancer and lung cancer). It was previously reported that plasma-derived sEVs from the healthy controls group were characterized by the absence of PD-L1 expression and that PD-L1 secreted in plasma sEVs from melanoma patients is useful to identify patients responding to immunotherapies ^24,48^. Importantly, we found that increase of PD-L1 levels was specific in melanoma patients with brain metastasis suggesting a strong immunosuppressive systemic change. It would be interesting to monitor PD-L1 in the plasma of melanoma patients with brain metastasis as a potential biomarker of response to therapy in this specific subtype. However, the study of plasma EVs is limited by the uncertainty of the cellular origin of these vesicles. Previous studies have suggested that most of the material found in blood would have an immunological origin ^49,50^. Unfortunately, in our study we could not identify the origin of STAT3 and PD-L1 in circulation, the identification of the source of these molecules would be important in future studies to determine whether they reflect systemic or tumor changes along metastatic progression.

Considering the role of EVs in the pre-metastatic niche conformation, the analysis of these vesicles using blood, the main route of neoplastic dissemination to the brain, could inform molecular features with potential to predict the risk of brain metastases. However, this study has several limitations. Despite the total number of the series exceeds 120 patients, the analyses according to the tumor subtypes are limited by a low sample size, which only allows the establishment of hypotheses for future studies. Nevertheless, the inclusion of different types of solid tumors favors the generalization of these results.

The samples were only collected at the time of the recent diagnosis of CNS dissemination; this fact makes it difficult the interpretation of the findings between true molecular risk factors predisposing to brain metastases or the systemic expression of already established brain lesions. Dynamic follow-up studies, including sample collections at different points during the clinical course (before and after the diagnosis of brain involvement) will be useful for the interpretation of the results here presented. Moreover, the intrinsic limitations of the imaging techniques currently used for the diagnosis could lead to a potential selection bias difficult to assess for in this series.

Finally, the complexity of cancer biology makes it difficult to explain oncological events, such as tumor dissemination to the brain, by means of a single approach. There are genetic, environmental or immunological factors, among others, not considered in this work, which could also influence the results presented. In addition to clinical studies, *in vitro* and *in vivo* analyses are also needed to explain these results and to adequately assess their clinical significance.

Overall, this study suggests that plasma-circulating EVs present different quantitative and qualitative characteristics depending on the presence or absence of neoplastic brain dissemination, especially in melanoma patients. This information highlights the potential usefulness of EVs for the development of new biomarkers that could improve the care of oncological patients; however, functional studies defining the role of these vesicles in CNS metastases should be performed for an adequate interpretation of the data.

## Acknowledgements

This work was funded by RETOS SAF2017-82924-R (AEI/10.13039/501100011033/FEDER-UE), Fundación Ramón Areces, Fundación Bancaria “la Caixa” (HR18-00256) and Fundación Científica AECC (LABAE19027PEIN). We are also grateful for the support of the Translational Network for the Clinical Application of Extracellular Vesicles (TeNTaCLES), RED2018-102411-T (AEI/10.13039/501100011033), the Ramón y Cajal Programme, the FERO Foundation. The CNIO, certified as a Severo Ochoa Excellence Centre, is supported by the Spanish government through the ISCIII.

## Author information

### Contributions

A.C-G., L.M.S., E.C.G., D.C. and G.dV. provided patient samples. A.C-G.., S.S-R., M.H-R. and H.P. performed the sEV isolations and Western Blot analyses. A.C-G., S.S-R., M.H-R. and H.P. performed immmunohistochemical analysis. A.C-G., M.H-R. G.dV. and H.P. contributed to the analysis of results. A.C-G., L.M.S., G.dV. and H.P. conceived the original hypothesis. S.S-R., M.H-R., and H.P. planned the experiments. A.C-G., M.H-R., G.dV. and H.P. wrote the manuscript. H.P. directed and supervised the work. All the authors contributed to and approved the final version of the manuscript.

## Ethics declarations

The authors have no conflict of interests.

## Figure legends

**Supplementary figure 1. Survival analysis and graphical representation using Kaplan Meier showing the cumulative survival probability of the metastatic patients included with different solid tumors according to the absence (CNS-) or presence (CNS+) of central nervous system (CNS) metastases. A**. Lung cancer patients. **B**. Breast cancer patients. **C**. Kidney cancer patients. **D**. Melanoma patients. ns: not significant. Differences were assessed using the Log-Rank test.

**Supplementary figure 2. Study of pSTAT3/STAT3 and PD-L1 expression by Western blot in plasma-circulating sEVs from patients with lung and breast cancer. A**. Western blot image of the expression of different proteins (indicated on the right) in sEVs from lung cancer patients according to the absence (No) or presence (Yes) of central nervous system (CNS) metastases. **B**. Western blot image of the expression of different proteins (indicated on the right) in sEVs from breast cancer patients according to the absence (No) or presence (Yes) of CNS metastases. Luminal: luminal phenotype (breast cancer). HER2^+^: HER2-overexpressing phenotype (breast cancer). Triple negative: triple negative phenotype (breast cancer).

**Supplementary figure 3. Study of pSTAT3/STAT3 and PD-L1 expression by Western blot in plasma sEVs from patients with kidney cancer and healthy/cured controls. A**. Western blot image of the expression of different proteins (indicated on the right) in sEVs from kidney cancer patients according to the absence (No) or presence (Yes) of central nervous system (CNS) metastases. **B**. Western blot image of the expression of different proteins (indicated on the right) in sEVs from healthy/cured controls.

**Supplementary Table 1. Clinical and pathological parameters collected in this series of patients**. Central nervous system (CNS). American Joint Committee on Cancer (AJCC). Epidermal growth factor receptor (EGFR). Anaplastic Lymphoma kinase (ALK). Proto-oncogene receptor tyrosine kinase 1 (ROS1). Human epidermal growth factor receptor 2 (HER2). V-raf murine sarcoma viral oncogene homolog B1 (BRAF). * T (tumor) N (nodes) M (metastases). Edge SB, et al., eds. AJCC Cancer Staging Manual. 7th ed. New York: Springer-Verlag: 2009. ^#^ Assessed according to techniques commonly used in clinical practice: For lung cancer (EGFR, ALK, ROS1): Polymerase chain reaction (PCR), Fluorescence *in situ* hybridization (FISH), Next Generation Sequencing (NGS). For breast cancer (estrogen receptor, progresterone receptor, HER2): immunohistochemical assessment, FISH. For melanoma (BRAF): PCR, NGS. ^γ^ Specific therapeutic schemes depending on tumor histology. Supportive therapies with no evidence of antitumoral activity (e.g. denosumab) were excluded. The following treatments were considered: For lung cancer: chemotherapy, immune checkpoint inhibitors, anti-EGFR therapy, anti-ALK therapy, anti-ROS1 therapy. For breast cancer: chemotherapy, hormone therapy, cell cycle inhibitors, phosphoinositol 3-kinase (PI3K)/AKT/mammalian target of rapamycin (mTOR) pathway inhibitors, anti-HER2 therapies, immune checkpoint inhibitors. For kidney cancer: antiangiogenic therapy, immune checkpoint inhibitors, mTOR inhibitors. For melanoma: chemotherapy, immune checkpoint inhibitors, anti-BRAF therapy +/−anti-Mitogen-activated protein kinase (MEK) therapy.

**Supplementary Table 2. Clinical characteristics of the metastatic patients with central nervous system (CNS) involvement included in the study at the time of sample collection**. Data collected from metastatic patients with confirmed CNS involvement are included. * All patients presented stage IV at the time of sample collection. Central nervous system (CNS). International Metastatic Renal Cell Carcinoma Database Consortium (IMDC). Epidermal growth factor receptor (EGFR). Anaplastic Lymphoma kinase (ALK). Human epidermal growth factor receptor 2 (HER2). V-raf murine sarcoma viral oncogene homolog B1 (BRAF).

**Supplementary Table 3. Clinical characteristics of the metastatic patients without central nervous system (CNS) involvement included in the study at the time of sample collection**. Data collected from metastatic patients without CNS involvement are included. * All patients presented stage IV at the time of sample collection. Central nervous system (CNS). International Metastatic Renal Cell Carcinoma Database Consortium (IMDC). Epidermal growth factor receptor (EGFR). Anaplastic Lymphoma kinase (ALK). Human epidermal growth factor receptor 2 (HER2). V-raf murine sarcoma viral oncogene homolog B1 (BRAF).

**Supplementary Table 4. Quantification of particle number and protein concentration in samples from healthy/cured controls and patients without brain metastases**.

**Supplementary Table 5. Quantification of particle number and protein concentration in samples from patients with brain metastases**.

