## Supplementary figures and images for "Characterization of plasma circulating small extracellular vesicles in patients with metastatic solid tumors and newly diagnosed brain metastasis"

### Supplemental Figure 1

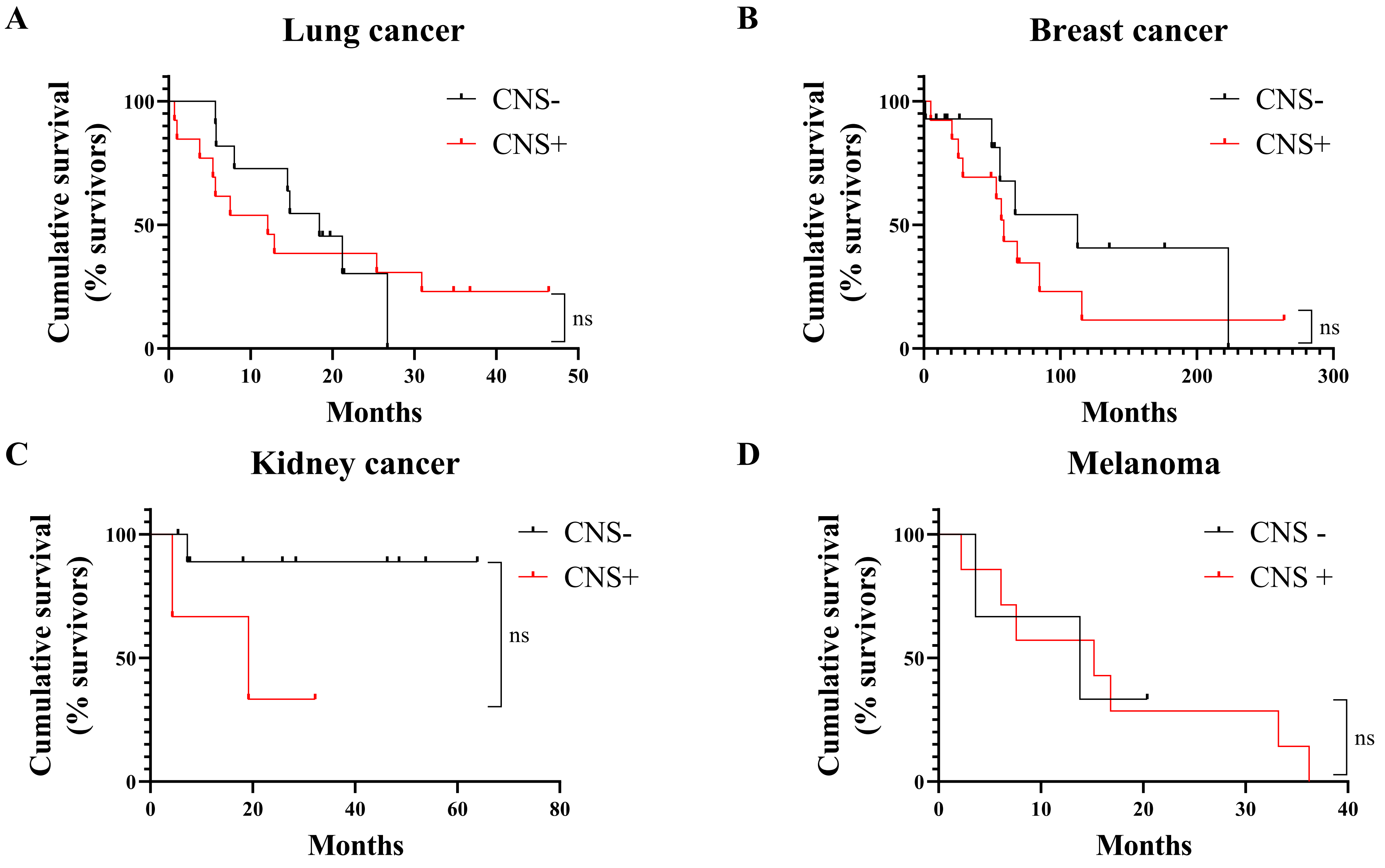


**Supplementary Figure 1**

### Supplemental Figure 2

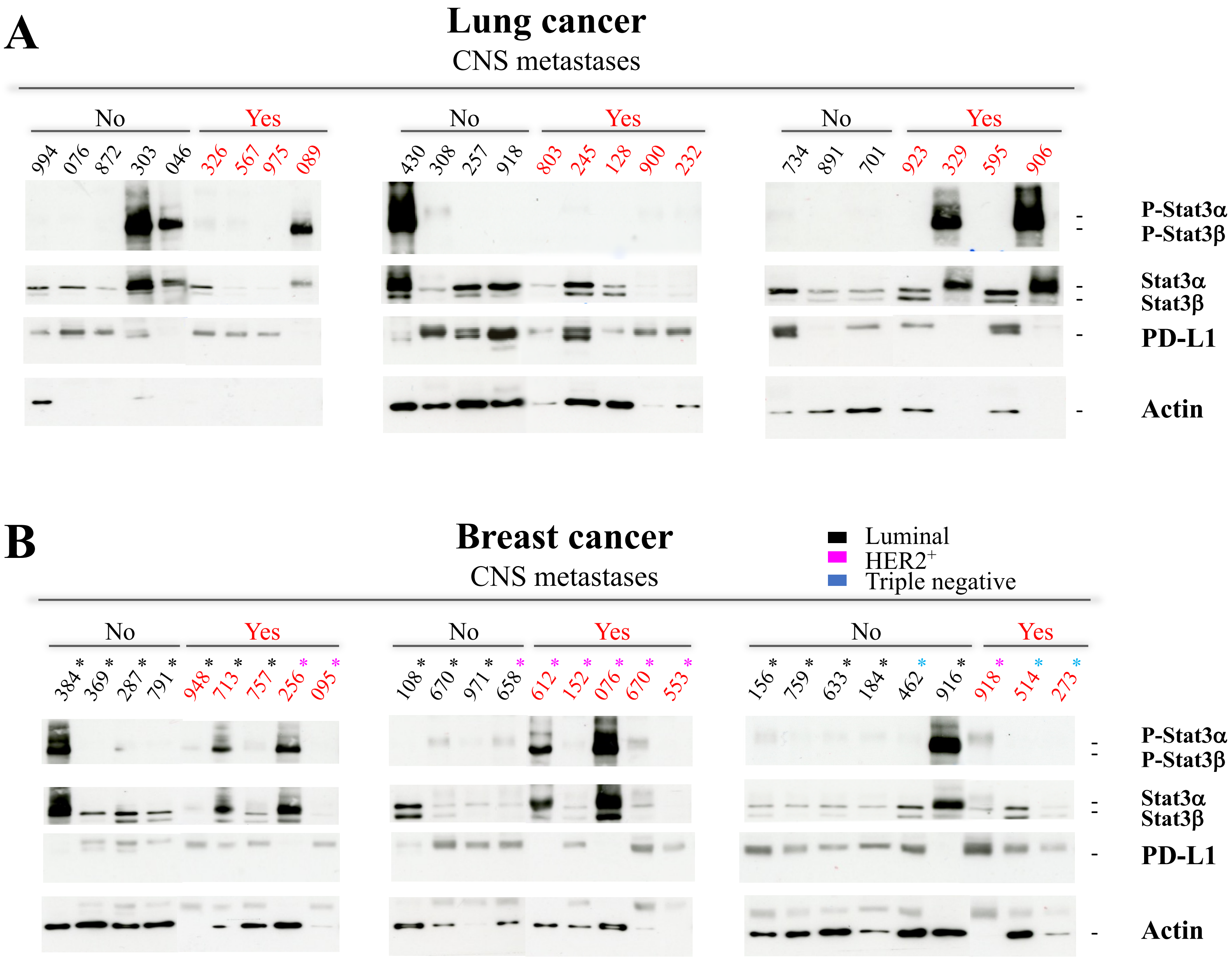


**Supplementary Figure 2**

### Supplemental Figure 3

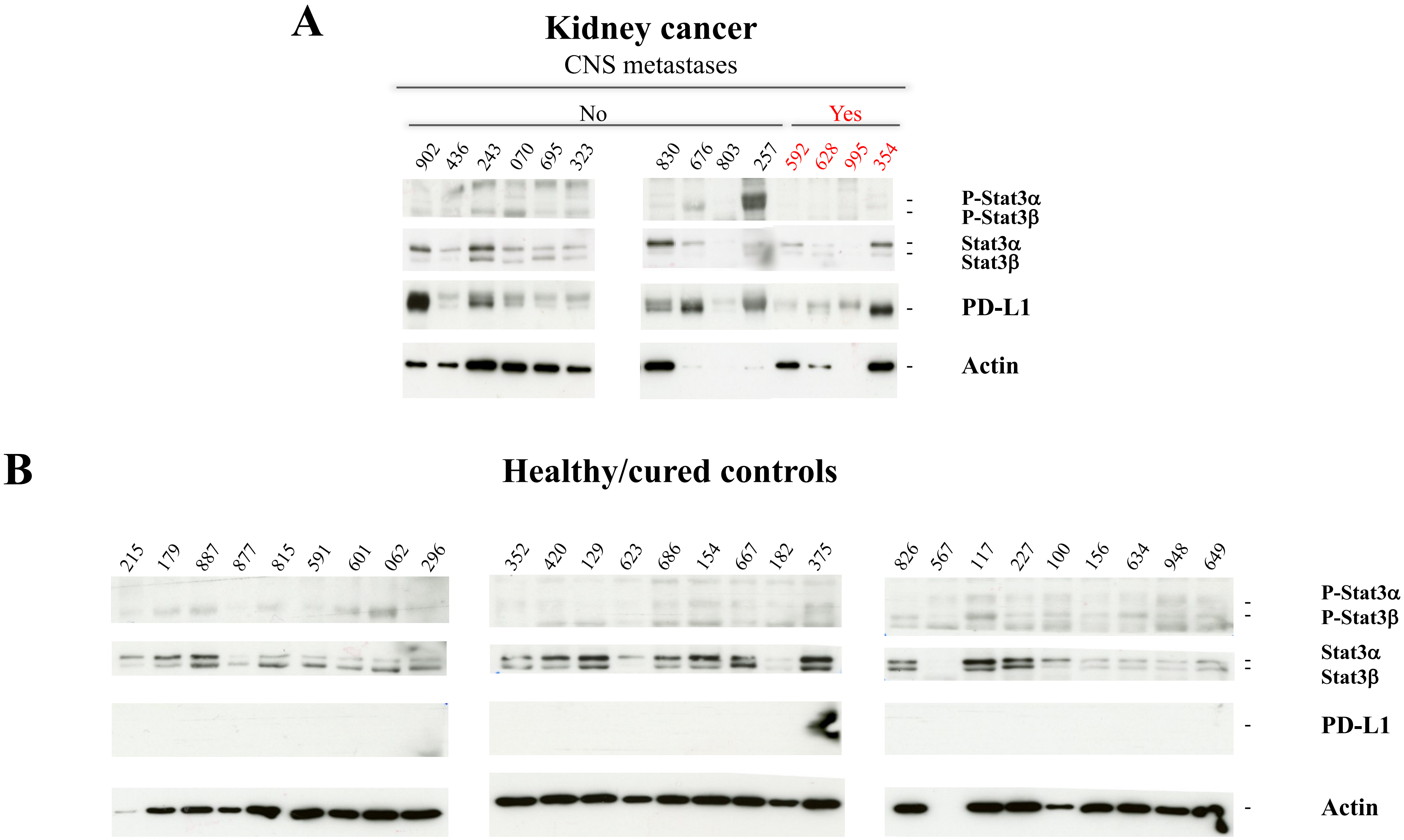


**Supplementary Figure 3**
