## Supplemental Table 1 for "Characterization of plasma circulating small extracellular vesicles in patients with metastatic solid tumors and newly diagnosed brain metastasis"

**Supplementary Table 1.** Clinical and pathological parameters collected in this series of patients

| **Patient characteristics** | |
| --- | --- |
| **Age**  (at the time of sample collection) | **Analytical values (blood count)**  (at the time of sample collection)  Leukocyte count (x1000/μl)  Neutrophil count (x1000/μl)  Lymphocyte count (x1000/μl) |
| **Gender**  Male  Female | **Date of death/last follow-up**  (as appropriate) |
| **Tumor characteristics** | |
| **Date of first diagnosis of the neoplastic disease**  (considered the date of anatomopathological diagnosis) | **Number of disease progressions (assessed clinically or radiologically) prior to sample collection**  0  1  2  3 or more |
| **Histological type of the primary tumor**  Lung cancer (non-small cell/small cell)  Breast cancer  Kidney cancer  Melanoma | **Type of tumor progression at the time of sample collection**  1: CNS without progression in other locations  2: CNS with progression in other locations  3: Other locations without CNS  4: No progression (healthy/cured controls) |
| **TNM stage at disease diagnosis**  (according to AJCC 7^th^ edition)*  I/II  III  IV | **Location of metastatic disease at the time of sample collection**  1: Bone/serosa only  2: Viscera (e.g. liver, lung)  3: CNS  1+2  1+3  2+3  1+2+3 |
| **Clinically relevant molecular alterations^#^**  For lung cancer: EGFR, ALK and ROS1 status  For breast cancer: Estrogen receptor, progesterone receptor and HER2 status  For melanoma: BRAF status | **Characteristics of recent CNS involvement**  New single lesion  2 or more lesions  Other (leptomeningeal carcinomatosis) |
| **Date of confirmed metastatic disease**  (date of first clinical or radiological evidence of distant dissemination) |  |
| **Treatment characteristics** | |
| **Radical treatment received for the primary tumor**  Surgery  Radiotherapy  Concomitant chemotherapy + radiotherapy  Other | **Active systemic therapies ^γ^ received for metastatic disease**  (specific therapeutic schemes depending on tumor histology, including prior prophylactic CNS treatment) |
| **Active systemic therapies^γ^ received for localized disease**  (specific therapeutic schemes depending on tumor histology) | **Characteristics of active systemic therapy^γ^ received immediately after sample collection**  Including:  Therapeutic scheme (depending on histology)  Start date  Best response achieved  Date of radiological evidence for best response achieved  End date |
| **Number of previous active systemic therapies^γ^ received for metastatic disease at the time of sample collection**  0  1  2  3 or more |  |

**Supplementary Table 1.** **Clinical and pathological parameters collected in this series of patients.** Central nervous system (CNS). American Joint Committee on Cancer (AJCC). Epidermal growth factor receptor (EGFR). Anaplastic Lymphoma kinase (ALK). Proto-oncogene receptor tyrosine kinase 1 (ROS1). Human epidermal growth factor receptor 2 (HER2). V-raf murine sarcoma viral oncogene homolog B1 (BRAF). * T (tumor) N (nodes) M (metastases). Edge SB, et al., eds. AJCC Cancer Staging Manual. 7th ed. New York: Springer-Verlag: 2009. ^#^ Assessed according to techniques commonly used in clinical practice: For lung cancer (EGFR, ALK, ROS1): Polymerase chain reaction (PCR), Fluorescence *in situ* hybridization (FISH), Next Generation Sequencing (NGS). For breast cancer (estrogen receptor, progresterone receptor, HER2): immunohistochemical assessment, FISH. For melanoma (BRAF): PCR, NGS. **^γ^** Specific therapeutic schemes depending on tumor histology. Supportive therapies with no evidence of antitumoral activity (e.g. denosumab) were excluded. The following treatments were considered: For lung cancer: chemotherapy, immune checkpoint inhibitors, anti-EGFR therapy, anti-ALK therapy, anti-ROS1 therapy. For breast cancer: chemotherapy, hormone therapy, cell cycle inhibitors, phosphoinositol 3-kinase (PI3K)/AKT/mammalian target of rapamycin (mTOR) pathway inhibitors, anti-HER2 therapies, immune checkpoint inhibitors. For kidney cancer: antiangiogenic therapy, immune checkpoint inhibitors, mTOR inhibitors. For melanoma: chemotherapy, immune checkpoint inhibitors, anti-BRAF therapy +/- anti-Mitogen-activated protein kinase (MEK) therapy.
