## Supplemental Table 2 for "Characterization of plasma circulating small extracellular vesicles in patients with metastatic solid tumors and newly diagnosed brain metastasis"

| **Supplementary Table 2.** Clinical characteristics of the metastatic patients with central nervous system (CNS) involvement included in the study at the time of sample collection | | | | | | | | | | | | |
| --- | --- | --- | --- | --- | --- | --- | --- | --- | --- | --- | --- | --- |
|  |  | **Lung cancer (n=13)** | |  | **Breast cancer (n=14)** | |  | **Kidney cancer (n=4)** | |  | **Melanoma (n=11)** | |
| **TNM stage at disease diagnosis** |  | **n** | **%** |  | **n** | **%** |  | **n** | **%** |  | **n** | **%** |
| I/ II |  | 0 | 0 |  | 6 | 43 |  | 1 | 25 |  | 2 | 18 |
| III |  | 2 | 15 |  | 3 | 21 |  | 0 | 0 |  | 4 | 36 |
| IV* |  | 11 | 85 |  | 5 | 36 |  | 3 | 75 |  | 5 | 46 |
| **Histological subtype (lung cancer)** |  |  |  |  |  |  |  |  |  |  |  |  |
| Non-small cell |  | 13 | 100 |  | _ | _ |  | _ | _ |  | _ | _ |
| Small cell |  | 0 | 0 |  | _ | _ |  | _ | _ |  | _ | _ |
| **IMDC score (kidney cancer)** |  |  |  |  |  |  |  |  |  |  |  |  |
| Good prognosis |  | _ | _ |  | _ | _ |  | 1 | 25 |  | _ | _ |
| Intermediate prognosis |  | _ | _ |  | _ | _ |  | 0 | 0 |  | _ | _ |
| Bad prognosis |  | _ | _ |  | _ | _ |  | 3 | 75 |  | _ | _ |
| **Clinically relevant molecular alterations** |  |  |  |  |  |  |  |  |  |  |  |  |
| EGFR alterations (lung cancer) |  | 0 | 0 |  | _ | _ |  | _ | _ |  | _ | _ |
| ALK alterations (lung cancer) |  | 1 | 7 |  | _ | _ |  | _ | _ |  | _ | _ |
| No alterations (lung cancer) |  | 12 | 93 |  | _ | _ |  | _ | _ |  | _ | _ |
| Luminal phenotype (breast cancer) |  | _ | _ |  | 3 | 21 |  | _ | _ |  | _ | _ |
| HER2^+^ phenotype (breast cancer) |  | _ | _ |  | 8 | 58 |  | _ | _ |  | _ | _ |
| Triple negative phenotype (breast cancer) |  | _ | _ |  | 3 | 21 |  | _ | _ |  | _ | _ |
| BRAF alterations (melanoma) |  | _ | _ |  | _ | _ |  | _ | _ |  | 6 | 54 |
| No alterations (melanoma) |  | _ | _ |  | _ | _ |  | _ | _ |  | 5 | 46 |
| **Number of previous systemic therapies received for metastatic disease** |  |  |  |  |  |  |  |  |  |  |  |  |
| 0 |  | 8 | 62 |  | 0 | 0 |  | 2 | 50 |  | 8 | 73 |
| 1 or more |  | 5 | 38 |  | 14 | 100 |  | 2 | 50 |  | 3 | 27 |
| **Type of systemic therapies received for metastatic disease** |  |  |  |  |  |  |  |  |  |  |  |  |
| Chemotherapy |  | 4 | 31 |  | 13 | 93 |  | _ | _ |  | 0 | 0 |
| Immune checkpoint inhibitors |  | 2 | 15 |  | 2 | 14 |  | 1 | 25 |  | 2 | 18 |
| Targeted therapy (including hormone therapy) |  | 0 | 0 |  | 12 | 86 |  | 3 | 75 |  | 2 | 18 |
| No systemic treatment |  | 8 | 62 |  | 0 | 0 |  | 2 | 50 |  | 8 | 73 |
| **Type of tumor progression at the time of sample collection** |  |  |  |  |  |  |  |  |  |  |  |  |
| CNS without progression in other locations |  | 7 | 54 |  | 5 | 36 |  | 1 | 25 |  | 2 | 18 |
| CNS with progression in other locations |  | 6 | 46 |  | 9 | 64 |  | 3 | 75 |  | 9 | 82 |
| **Location of metastatic disease at the time of sample collection** |  |  |  |  |  |  |  |  |  |  |  |  |
| CNS only |  | 3 | 22 |  | 0 | 0 |  | 0 | 0 |  | 0 | 0 |
| CNS with bone/serosa |  | 0 | 0 |  | 0 | 0 |  | 0 | 0 |  | 0 | 0 |
| CNS with viscera |  | 5 | 39 |  | 5 | 36 |  | 1 | 25 |  | 9 | 82 |
| CNS with bone/serosa and viscera |  | 5 | 39 |  | 9 | 64 |  | 3 | 75 |  | 2 | 18 |
| **Characteristics of recent CNS involvement** |  |  |  |  |  |  |  |  |  |  |  |  |
| New single lesion |  | 5 | 39 |  | 2 | 14 |  | 2 | 50 |  | 3 | 27 |
| 2 or more lesions |  | 7 | 54 |  | 10 | 72 |  | 2 | 50 |  | 8 | 73 |
| Other (leptomeningeal carcinomatosis) |  | 1 | 7 |  | 2 | 14 |  | 0 | 0 |  | 0 | 0 |

**Supplementary Table 2.** **Clinical characteristics of the metastatic patients with central nervous system (CNS) involvement included in the study at the time of sample collection.** Data collected from metastatic patients with confirmed CNS involvement are included. * All patients presented stage IV at the time of sample collection. Central nervous system (CNS). International Metastatic Renal Cell Carcinoma Database Consortium (IMDC). Epidermal growth factor receptor (EGFR). Anaplastic Lymphoma kinase (ALK). Human epidermal growth factor receptor 2 (HER2). V-raf murine sarcoma viral oncogene homolog B1 (BRAF).
