## Supplemental Table 3 for "Characterization of plasma circulating small extracellular vesicles in patients with metastatic solid tumors and newly diagnosed brain metastasis"

| **Supplementary Table 3.** Clinical characteristics of the metastatic patients without central nervous system (CNS) involvement included in the study at the time of sample collection | | | | | | | | | | | | |
| --- | --- | --- | --- | --- | --- | --- | --- | --- | --- | --- | --- | --- |
|  |  | **Lung cancer (n=12)** | |  | **Breast cancer (n=21)** | |  | **Kidney cancer (n=10)** | |  | **Melanoma (n=7)** | |
| **TNM stage at disease diagnosis** |  | **n** | **%** |  | **n** | **%** |  | **n** | **%** |  | **n** | **%** |
| I/ II |  | 0 | 0 |  | 11 | 53 |  | 3 | 30 |  | 3 | 42 |
| III |  | 2 | 17 |  | 6 | 27 |  | 2 | 20 |  | 2 | 29 |
| IV* |  | 10 | 83 |  | 4 | 20 |  | 5 | 50 |  | 2 | 29 |
| **Histological subtype (lung cancer)** |  |  |  |  |  |  |  |  |  |  |  |  |
| Non-small cell |  | 10 | 83 |  | _ | _ |  | _ | _ |  | _ | _ |
| Small cell |  | 2 | 17 |  | _ | _ |  | _ | _ |  | _ | _ |
| **IMDC score (kidney cancer)** |  |  |  |  |  |  |  |  |  |  |  |  |
| Good prognosis |  | _ | _ |  | _ | _ |  | 2 | 20 |  | _ | _ |
| Intermediate prognosis |  | _ | _ |  | _ | _ |  | 6 | 60 |  | _ | _ |
| Bad prognosis |  | _ | _ |  | _ | _ |  | 2 | 20 |  | _ | _ |
| **Clinically relevant molecular alterations** |  |  |  |  |  |  |  |  |  |  |  |  |
| EGFR alterations (lung cancer) |  | 1 | 8 |  | _ | _ |  | _ | _ |  | _ | _ |
| ALK alterations (lung cancer) |  | 0 | 0 |  | _ | _ |  | _ | _ |  | _ | _ |
| No alterations (lung cancer) |  | 11 | 92 |  | _ | _ |  | _ | _ |  | _ | _ |
| Luminal phenotype (breast cancer) |  | _ | _ |  | 17 | 80 |  | _ | _ |  | _ | _ |
| HER2^+^ phenotype (breast cancer) |  | _ | _ |  | 2 | 10 |  | _ | _ |  | _ | _ |
| Triple negative phenotype (breast cancer) |  | _ | _ |  | 2 | 10 |  | _ | _ |  | _ | _ |
| BRAF alterations (melanoma) |  | _ | _ |  | _ | _ |  | _ | _ |  | 3 | 42 |
| No alterations (melanoma) |  | _ | _ |  | _ | _ |  | _ | _ |  | 4 | 58 |
| **Number of previous systemic therapies received for metastatic disease** |  |  |  |  |  |  |  |  |  |  |  |  |
| 0 |  | 9 | 75 |  | 9 | 43 |  | 4 | 40 |  | 6 | 87 |
| 1 or more |  | 3 | 25 |  | 12 | 57 |  | 6 | 60 |  | 1 | 13 |
| **Type of systemic therapies received for metastatic disease** |  |  |  |  |  |  |  |  |  |  |  |  |
| Chemotherapy |  | 3 | 25 |  | 8 | 38 |  | _ | _ |  | 0 | 0 |
| Immune checkpoint inhibitors |  | 0 | 0 |  | 2 | 10 |  | 1 | 10 |  | 1 | 13 |
| Targeted therapy (including hormone therapy) |  | 0 | 0 |  | 18 | 86 |  | 7 | 70 |  | 0 | 0 |
| No systemic treatment |  | 9 | 75 |  | 9 | 43 |  | 4 | 40 |  | 6 | 87 |
| **Location of metastatic disease at the time of sample collection** |  |  |  |  |  |  |  |  |  |  |  |  |
| Bone/serosa only |  | 0 | 0 |  | 1 | 5 |  | 2 | 20 |  | 1 | 13 |
| Viscera only |  | 5 | 42 |  | 3 | 15 |  | 3 | 30 |  | 4 | 58 |
| Bone/serosa with viscera |  | 7 | 58 |  | 17 | 80 |  | 5 | 50 |  | 2 | 29 |

**Supplementary Table 3.** **Clinical characteristics of the metastatic patients without central nervous system (CNS) involvement included in the study at the time of sample collection.** Data collected from metastatic patients without CNS involvement are included. * All patients presented stage IV at the time of sample collection. Central nervous system (CNS). International Metastatic Renal Cell Carcinoma Database Consortium (IMDC). Epidermal growth factor receptor (EGFR). Anaplastic Lymphoma kinase (ALK). Human epidermal growth factor receptor 2 (HER2). V-raf murine sarcoma viral oncogene homolog B1 (BRAF).
