## Supplemental Table 4 for "Characterization of plasma circulating small extracellular vesicles in patients with metastatic solid tumors and newly diagnosed brain metastasis"

**Supplementary Table 4.** Quantification of particle number and protein concentration in samples from healthy/cured controls and patients without brain metastases

| **Healthy/cured controls and patients without brain metastases** | | | | | | | | | | | | | | |
| --- | --- | --- | --- | --- | --- | --- | --- | --- | --- | --- | --- | --- | --- | --- |
| Healthy/cured controls | | | Lung cancer | | | Breast cancer | | | Kidney cancer | | | Melanoma | | |
| # | Particles (x10^8^)/ml | Protein (µg)/ml | # | Particles (x10^8^)/ml | Protein (µg)/ml | # | Particles (x10^8^)/ml | Protein (µg)/ml | # | Particles (x10^8^)/ml | Protein (µg)/ml | # | Particles (x10^8^)/ml | Protein (µg)/ml |
| 215 | 1.9 | 7.07 | 994 | 6.2 | 23.6 | 582 | 0.9 | 2.29 | 830 | 0.97 | 3.69 | 918 | 2.9 | 1.46 |
| 149 | 1.5 | 5.93 | 76 | 3.3 | 12.51 | 377 | 3.1 | 2.34 | 676 | 2.1 | 12 | 628 | 15 | 37.67 |
| 179 | 2.3 | 4.9 | 872 | 2.2 | 35.17 | 762 | 1 | 8.42 | 803 | 23 | 16.48 | 855 | 2.7 | 17.69 |
| 192 | 2.1 | 0.56 | 303 | 0.88 | 10.95 | 916 | 9.7 | 5.53 | 257 | 1.6 | 86.95 | 343 | 3.1 | 31.82 |
| 887 | 1.9 | 5.42 | 46 | 3.2 | 10.81 | 448 | 2 | 2.76 | 902 | 2.7 | 6.21 | 864 | 0.92 | 1.2 |
| 877 | 2.5 | 5.87 | 430 | 5.8 | 52.36 | 384 | 5.5 | 17.63 | 436 | 2.2 | 25.6 | 392 | 3.7 | 17.55 |
| 815 | 6.6 | 3.92 | 308 | 1.6 | 21.08 | 369 | 4 | 3.96 | 243 | 5.1 | 12.08 | 825 | 11 | 56.67 |
| 591 | 6.9 | 4.51 | 257 | 2.6 | 14.14 | 287 | 5.6 | 8.48 | 70 | 5.7 | 34.85 |  |  |  |
| 626 | 1.6 | 2.54 | 918 | 8.5 | 4.93 | 791 | 2.7 | 5.22 | 695 | 3.3 | 20.2 |  |  |  |
| 601 | 2.6 | 7.63 | 734 | 13 | 21.47 | 816 | 3.1 | 0.6 | 323 | 1.7 | 29.97 |  |  |  |
| 875 | 19 | 3.09 | 891 | 3.5 | 2.77 | 108 | 7.2 | 4.19 |  |  |  |  |  |  |
| 62 | 5.9 | 13.93 | 701 | 2.2 | 3.34 | 670 | 0.83 | 11.95 |  |  |  |  |  |  |
| 296 | 17 | 4.6 |  |  |  | 971 | 2.5 | 22.04 |  |  |  |  |  |  |
| 352 | 6.6 | 15.37 |  |  |  | 658 | 2.2 | 30.97 |  |  |  |  |  |  |
| 420 | 1.9 | 12.32 |  |  |  | 156 | 5.1 | 45.16 |  |  |  |  |  |  |
| 129 | 6.3 | 15.42 |  |  |  | 759 | 4.5 | 41.01 |  |  |  |  |  |  |
| 623 | 2 | 23.1 |  |  |  | 633 | 1.6 | 43.93 |  |  |  |  |  |  |
| 686 | 2.8 | 17.1 |  |  |  | 184 | 1.3 | 38.83 |  |  |  |  |  |  |
| 154 | 2.1 | 8.91 |  |  |  | 462 | 6.5 | 24.78 |  |  |  |  |  |  |
| 667 | 2.6 | 19.33 |  |  |  | 350 | 6.5 | 40.5 |  |  |  |  |  |  |
| 182 | 3.1 | 15.78 |  |  |  | 378 | 5.7 | 32.3 |  |  |  |  |  |  |
| 375 | 3.8 | 22.57 |  |  |  |  |  |  |  |  |  |  |  |  |
| 826 | 11 | 19.78 |  |  |  |  |  |  |  |  |  |  |  |  |
| 567 | 0.52 | 76.45 |  |  |  |  |  |  |  |  |  |  |  |  |
| 117 | 1.5 | 19.07 |  |  |  |  |  |  |  |  |  |  |  |  |
| 227 | 2.5 | 17.08 |  |  |  |  |  |  |  |  |  |  |  |  |
| 100 | 4.1 | 36.07 |  |  |  |  |  |  |  |  |  |  |  |  |
| 156 | 4.3 | 22.15 |  |  |  |  |  |  |  |  |  |  |  |  |
| 634 | 3.3 | 30.77 |  |  |  |  |  |  |  |  |  |  |  |  |
| 948 | 3.6 | 31.68 |  |  |  |  |  |  |  |  |  |  |  |  |
| 649 | 3.8 | 17.45 |  |  |  |  |  |  |  |  |  |  |  |  |
