## Supplemental Table 5 for "Characterization of plasma circulating small extracellular vesicles in patients with metastatic solid tumors and newly diagnosed brain metastasis"

**Supplementary Table 5.** Quantification of particle number and protein concentration in samples from patients with brain metastases

| **Patients with brain metastases** | | | | | | | | | | | |
| --- | --- | --- | --- | --- | --- | --- | --- | --- | --- | --- | --- |
| Lung cancer | | | Breast cancer | | | Kidney cancer | | | Melanoma | | |
| # | Particles (x10^8^)/ml | Protein (µg)/ml | # | Particles (x10^8^)/ml | Protein (µg)/ml | # | Particles (x10^8^)/ml | Protein (µg)/ml | # | Particles (x10^8^)/ml | Protein (µg)/ml |
| 326 | 2 | 26.72 | 948 | 0.6 | 164.04 | 592 | 0.79 | 20.32 | 687 | 1.2 | 10.6 |
| 567 | 0.73 | 52.51 | 713 | 1 | 79.17 | 628 | 0.79 | 87.05 | 31 | 1.4 | 86.73 |
| 975 | 0.84 | 42.14 | 757 | 2.3 | 17.37 | 995 | 1.1 | 214.2 | 641 | 1.6 | 12.02 |
| 89 | 1.2 | 85.1 | 256 | 0.53 | 21.72 | 354 | 7 | 11.91 | 993 | 3.6 | 37.47 |
| 803 | 0.95 | 55.62 | 95 | 1.6 | 23.73 |  |  |  | 64 | 4.7 | 19.15 |
| 245 | 1.3 | 1.83 | 612 | 3.5 | 17.46 |  |  |  | 140 | 0.75 | 3.12 |
| 128 | 1.6 | 30.57 | 152 | 2.2 | 50.37 |  |  |  | 901 | 2.4 | 13.41 |
| 900 | 0.55 | 52.1 | 76 | 3.1 | 16.77 |  |  |  | 456 | 1.3 | 12.02 |
| 232 | 1.2 | 38.26 | 670 | 1 | 45.98 |  |  |  | 613 | 2.6 | 14.26 |
| 923 | 3.7 | 22.19 | 553 | 1.5 | 113.18 |  |  |  | 960 | 11 | 15.91 |
| 329 | 1.1 | 26.11 | 918 | 1.7 | 34.02 |  |  |  | 77 | 23 | 43.01 |
| 595 | 4 | 21.56 | 514 | 3.7 | 51.73 |  |  |  |  |  |  |
| 906 | 3.2 | 97.49 | 273 | 0.61 | 13.22 |  |  |  |  |  |  |
|  |  |  | 563 | 0,.9 | 17.61 |  |  |  |  |  |  |
